# Livestock and microcephaly, traces of an association?

**DOI:** 10.1101/087825

**Authors:** Ion de Andrade

## Abstract

While there is no doubt about the participation of Zika virus in microcephaly, its epidemiology is not entirely clear and doubts remain about the intervention of other factors. In studies on the epidemiology of dengue, the infestation by Aedes aegypti peridomiciliary and the population density are the main determinants for viral spread. However, in Rio Grande do Norte state (RN), the counties that have confirmed cases of microcephaly overlapped the river basins regions surrounded by agriculture and livestock. In addition, the prevalence of microcephaly at the end of the first year of the epidemic was higher in small towns than in larger ones, elements that seem to contradict what is known about the epidemic by other arboviruses. Methods: 234 cases of microcephaly were analyzed from three states and 144 counties. Results: An exponential trend of higher prevalence of microcephaly in the smaller cities (r^2^=0,7121) was found.The correlation coefficients (R) between the Prevalence of microcephaly and the variables that measured the density of animals in the territory ranged from moderate to strong. Discussion: Concerning microcephaly, studies in progress point to the possibility of association between the Zika Virus and the BVDV, a virus known to produce birth defects in farm animals but perceived as innocuous in humans. Conclusions: The overlap of cases of microcephaly in river basins, their higher prevalence in smaller cities, the strength of the correlation coefficient, render necessary new etiological and pathophysiological studies.

**ABBREVIATIONS:**

BVDV: Bovine diarrhea virus
CE: Ceará state
IBGE: Instituto Brasileiro of Geography and Statistics
IPESQ: Instituto de Pesquisa da Paraíba
PB: Paraíba state
RN: Rio Grande do Norte state
UFRJ: Rio de Janeiro Federal University
ZKV: Zika Virus.

## 6. INTRODUCTION

With 2033 confirmed cases of microcephaly, Brazil is the main country affected worldwide [1].

Although studies have identified the Zika Virus (ZKV) in fetal tissues and in babies with microcephaly [2-5] its epidemiology is still not fully understood. It is not clear, for instance, why only the smallest part of mothers who had Zika in the first trimester of pregnancy gave birth to babies affected by this complication [6-8]. Brazil is also the country where microcephaly is more frequent considering the reported cases of Zika [9, 10]. The predominance of microcephaly in the Northeastern region, its higher prevalence in small towns and its greater frequency in low-income children, points to other possible intervening factors [7, 8] and begins to create doubts about the genesis of the problems as an isolated action of the ZKV, showing the epidemic is not entirely resolved concerning the etiological point of view [11, 12].

Studies conducted by the Federal University of Rio de Janeiro (UFRJ) and Paraiba Research Institute (IPESQ) [12] are considering the possibility that the ZKV can act as a facilitator of the infection by the BVDV [13], a virus known to cause malformations in cattle and pigs, but considered incapable, under normal circumstances, to produce disease in humans [14]. Although fragments of this virus have been found in fetal tissues from microcephalic babies, BVDV is known to contaminate vaccines and other biological materials used in human healthcare, which makes this finding still inconclusive [14].

This study brings new elements to the hypothesis of livestock participation in the epidemiology of microcephaly, adding one more element to the thesis of involvement of the BVDV.

## 7. METHODOLOGY

### Population

Concerning the analysis it was considered: a) the cases of microcephaly confirmed by the state health secretariats of Rio Grande do Norte (RN), Ceará (CE) and Paraíba (PB), available at the time of data analysis [15-17]; b) the population of cattle and pigs, according to the 2006 IBGE agricultural census of Rio Grande do Norte, Ceará and Paraíba [18], c) the population by county (RN, CE and PB) according to the projection of the IBGE for 2012 [19]; d) the number of live births by county for 2014, according to MS / SINASC [20] and d) the area in Km^2^ of the counties (RN, CE and PB) according to the IBGE [21]. The latest available data from each serie was used for the analysis.

The cases of microcephaly coming from counties with more than 100,000 inhabitants were excluded from this analysis due to the lower interface of this population (more urban) with livestock, and because these counties are regional health centers for a micro-region and receive from there numerous patients, compromising the statistical reliability of their data. Therefore, regarding this: a) in Rio Grande do Norte state, the counties of Natal, Mossoró and Parnamirim 58 cases out of 135 were excluded; b) in Ceará state, the counties of Fortaleza, Caucaia, Juazeiro, Maracanaú, Sobral, Crato and Maranguape, totaling 65 cases out of 136; and c) Paraíba state the counties of João Pessoa, Campina Grande, Santa Rita, Patos and Bayeux, resulting in 62 cases out of 148.

The sample included 77 confirmed cases from 44 counties of RN state; 71 confirmed cases from 45 counties of Ceará state and 86 confirmed cases from 55 counties of Paraiba state, totaling 234 cases out of 419; a sufficient sample for analysis with a 95% confidence level, assuming an average error margin of 5% for all the data.

### Variables

Five variables, three population-based and two territory-based were created. The population-based variables divided the population of bovines and swine by the counties population. Because these variables followed a similar formulation, they were all called “Prevalence”. These were the ***Prevalence of swine*** and the ***Prevalence of bovine***. The variable ***Prevalence of Microcephaly*** was calculated with live birth population. The variable ***County Population*** measured the population of counties from RN, Ceará and Paraíba and was imported from IBGE. It served to graphically distribute Prevalence of Microcephaly.

The territorial-based variables divided the prevalence of swine and bovine by the areas of the counties, creating "densities". These variables received the generic name of "densities of prevalence". These were the ***Density of Prevalence of swine*** and the ***Density of Prevalence of bovines***.

### Analysis

The variables that measured the prevalence were calculated using the following formulas: ***Prevalence of microcephaly*** [(cases of microcephaly / live births) x1000]; ***Prevalence of Bovines*** [(Bovine population / county population) x1000] and ***Prevalence of Swine*** [(Swine population / county population) x1000]. The variables that measured the density of the prevalence of bovines and swine were built by dividing these previously calculated prevalences by the area of each county.The resulting ratios were multiplied by a thousand.

The variable ***Prevalence of microcephaly*** was submitted to the linear regression with the variables ***Prevalence of Swine***, ***Prevalence of Bovines***, ***Density of Prevalence of Swine*** and ***Density of Prevalence of Bovines***. Each variable was applied to the states of Rio Grande do Norte, Ceará and Paraíba and to the total of combined cases. The analysis included regressions by state and total.

The RN, CE and PB acronyms distinguished the variables by state. Confirmed cases of microcephaly were plotted on the municipal map of Rio Grande do Norte state with TABWIN 32.

The hydrogeological map of Rio Grande do Norte state was imported from the database of the National Water Agency [22, 23] and the map of risk for dengue epidemic was imported from the Secretariat of Public Health of RN state [24]. The spatial distribution of dengue cases from the National System of notifiable diseases (SINAN dengue online) was plotted on the map of Rio Grande do Norte state [25] with TABWIN 32.

## 8. RESULTS

The Prevalence of microcephaly was higher in smaller counties, a finding that highlights a rurality of the disease that should not occur. Epidemics transmitted by the Aedes aegypti are conditioned by the peridomiciliary infestation gradient and by the population density. The prevalence of microcephaly should therefore be associated with these conditions.

The spatial distribution of the prevalence of microcephaly (Figure 2) overlapped the river basins of the state (Figure 3) countering again the logic of spreading by the gradient of peridomiciliary infestation and by the population density.

**Figure 1.**
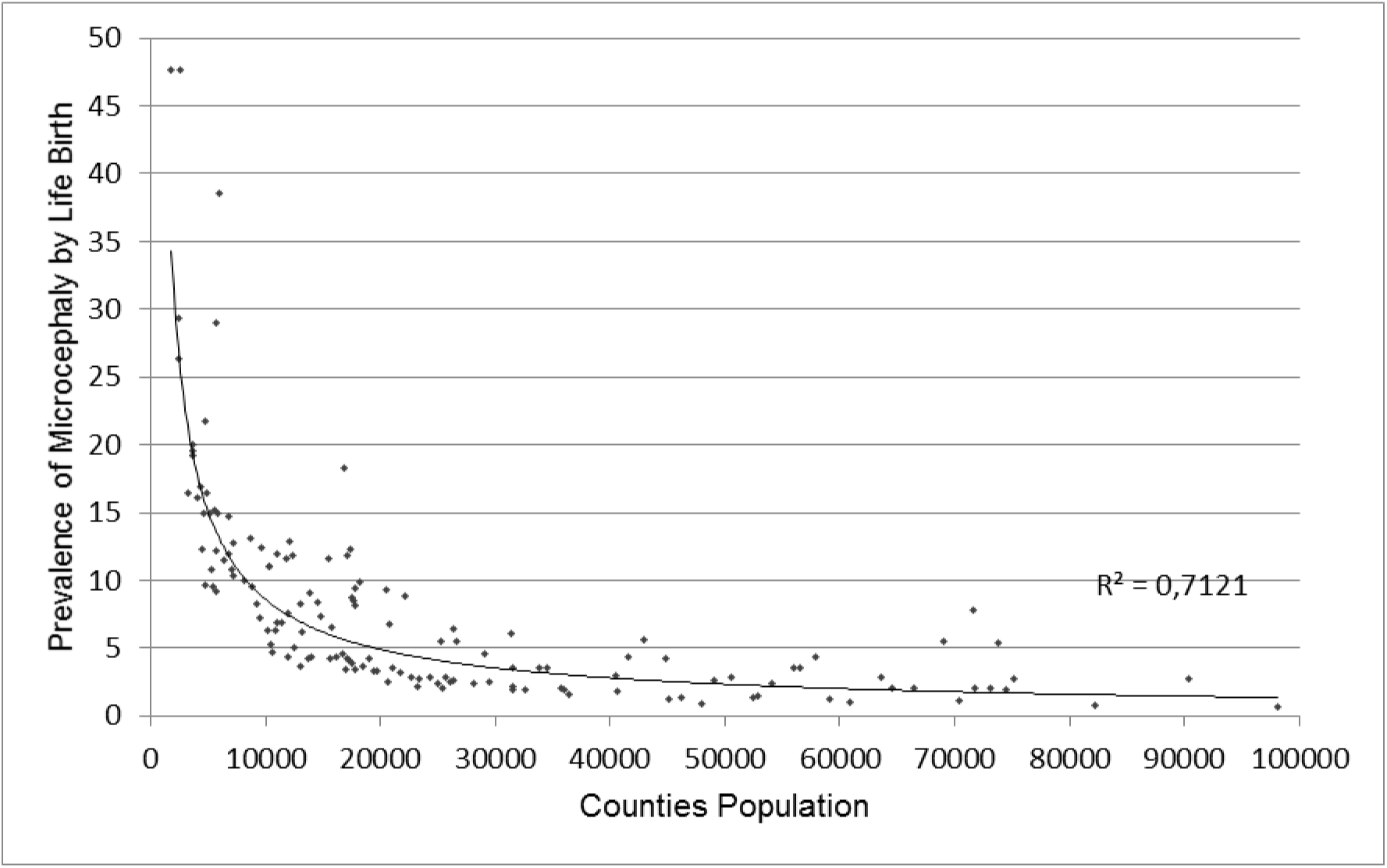
Distribution of the Prevalence of microcephaly by Live Births in the counties of RN, CE and PB according to the county population.

**Figure 2.**
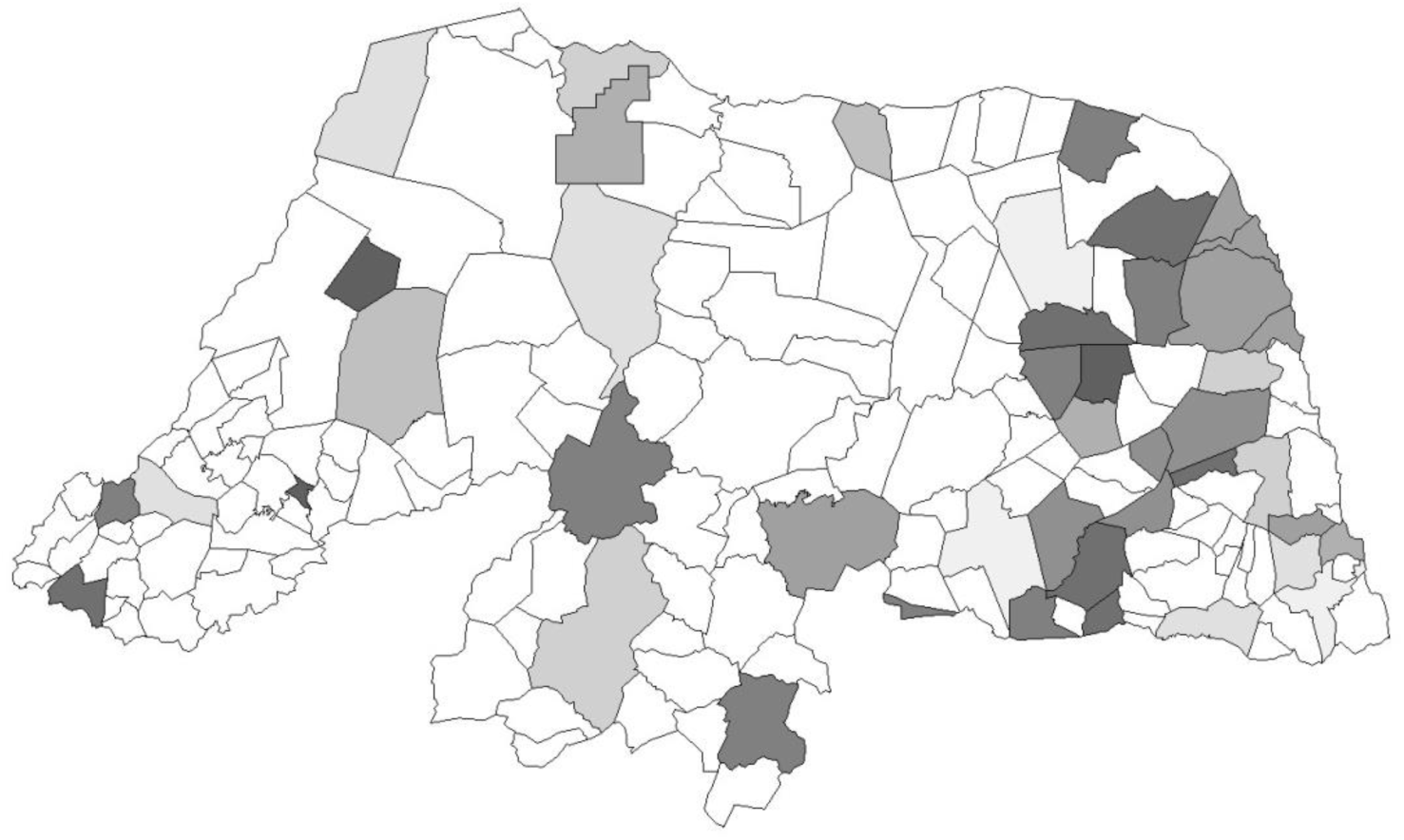
Spatial distribution of the Prevalence of microcephaly by Live Births in the counties of Rio Grande do Norte. (RIO GRANDE DO NORTE, SESAP-RN, Epidemiological Bulletin, Week 32, 2016)

Added to this the fact that the central region of the state, where there were registered high levels of infestation by Aedes aegypti, (Figure 4) did not have registered cases of microcephaly (Figure 2). On the other hand, in the north-south axis, which followed the river Piranhas-Açu (Figure 3), where registered infestation rates were low (Figure 4), there have been several reported cases (Figure 2).

**Figure 3.**
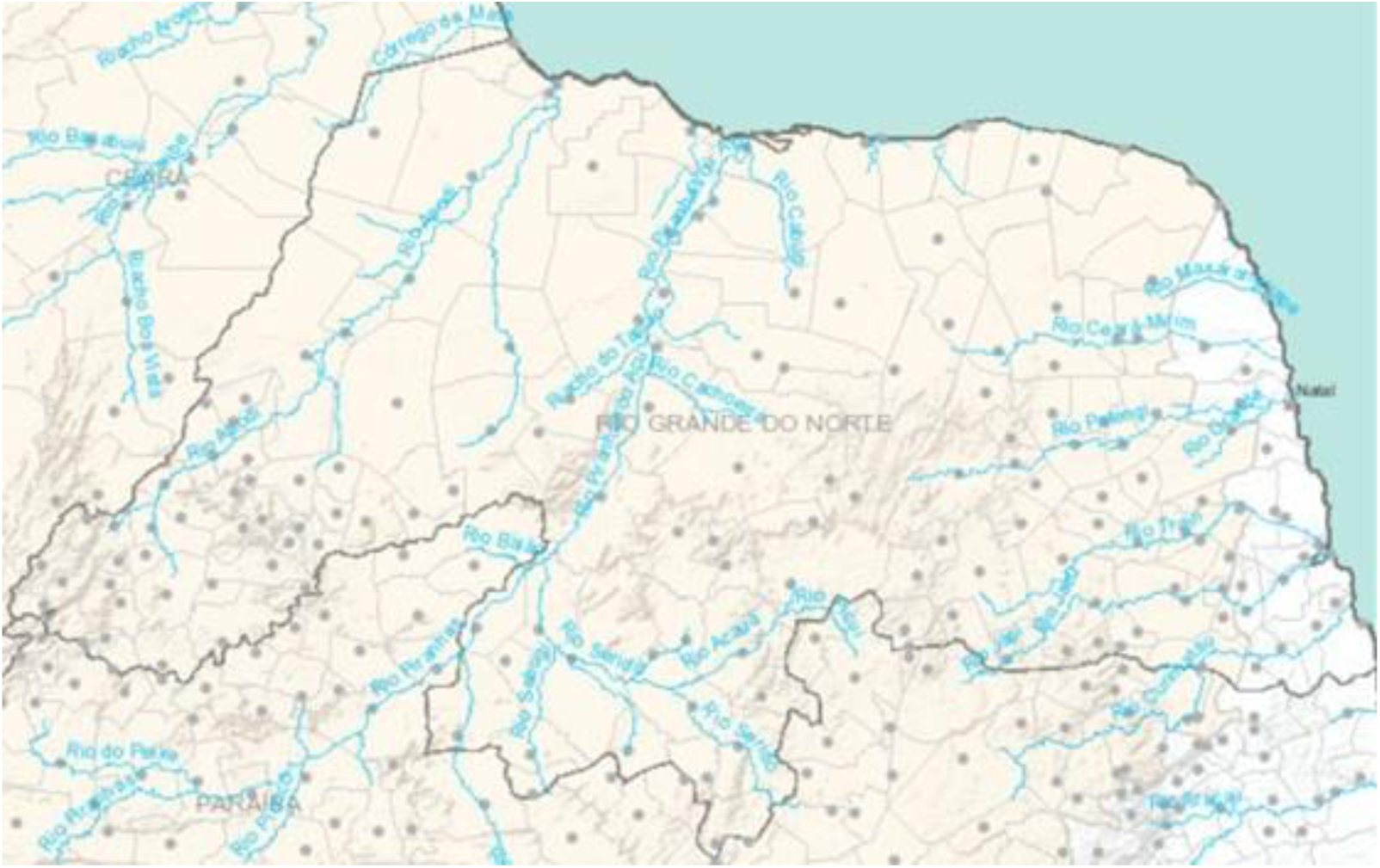
Rivers basin of Rio Grande do Norte state (BRASIL, Agência Nacional das Águas, Sistema de Monitoramento Hidrológico, 2016)

**Figure 4.**
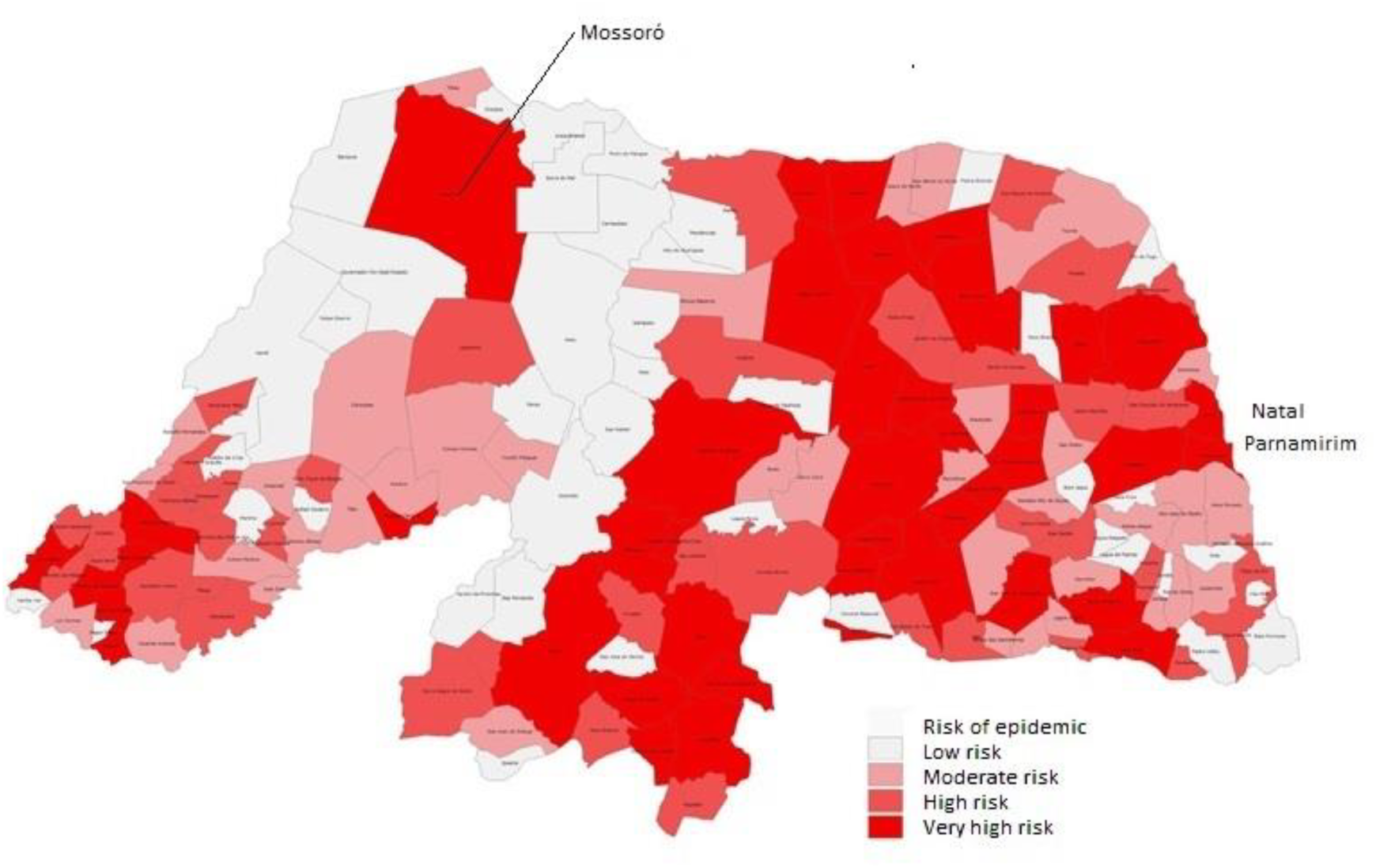
Spatial distribution of the risk of dengue epidemics for Rio Grande do Norte in 2014 (Rio Grande do Norte, SESAP RN, Mapa de Vulnerabilidade para a ocorrência de dengue no RN 2014)

The three counties of the state with populations over 100,000 (Natal, Mossoro and Parnamirim, marked on the map), excluded from the analysis, as explained in the methodology, appeared in 2014 as very high-risk areas for epidemic dengue due to the high levels of infestation by Aedes aegypti. However these counties, where the ideal conditions for the spread of ZKV were gathered, had a prevalence of microcephaly lower than the average of the counties with populations of less than 100,000 (2.9‰ against 4.9‰) countering the epidemiological logic.

The cases of dengue recorded in 2015 by the Information System of notifiable diseases, the SINAN (Figure 5) established higher prevalence in areas consistent with the risk assessed in 2014 (Figure 4). Although the risk assessed in 2014 and the 2015 epidemic have arisen from the peridomiciliary infestation indices by Aedes aegypti, a risk factor for all arboviruses and not only for dengue, there was no overlap of the risk (Figure 4) or of the epidemic (Figure 5) within the areas where cases of microcephaly were reporded (Figure 2).

**Figure 5.**
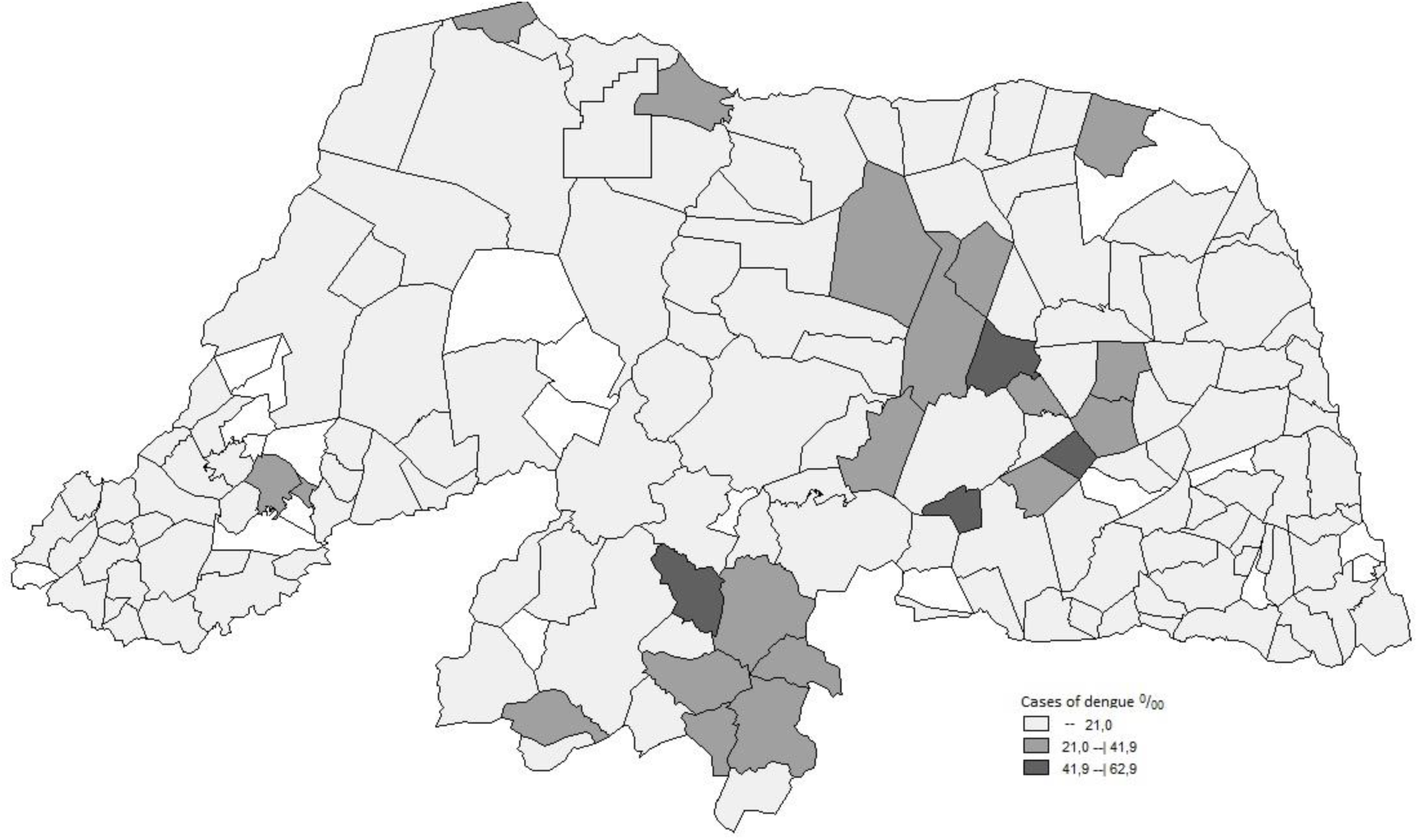
Spatial distribution of the dengue cases in 2015 according to the information system of notifiable diseases SINAN (BRASIL, Ministério da Saúde, DATASUS, SINAN dengue online)

The correlation coefficients found between the variables that measured the prevalence of bovines and swine and Prevalence of microcephaly (Table 1) were insufficient and inconclusive.

**Table 1 -.** Correlation coefficient (R) between the Prevalence of microcephaly (by Live Births) and the Prevalence of Swine and Bovines for the counties of Rio Grande do Norte, Ceará and Paraíba.

| Variável | R | p value |
| --- | --- | --- |
| *Prevalence of Swine RN* | 0,6314 | < 0,0001 |
| *Prevalence of Bovines RN* | 0,4205 | < 0,05 |
| *Prevalence of Swine CE* | 0,0129 | 0,9326 |
| *Prevalence of Bovines CE* | 0,2425 | 0,1083 |
| *Prevalence of Swine PB* | 0,3258 | < 0,05 |
| *Prevalence of Bovines PB* | 0,3087 | < 0,05 |
| *Prevalence of Swine (RN, PB e CE)* | 0,0012 | 0,9883 |
| *Prevalence of Bovines (RN, PB e CE)* | 0,3932 | < 0,0001 |

However, the inclusion of the area of the counties in the calculation of the variables allowed us to consider the density of animals in the territory, which sensitized the calculation, homogenized their results and increased correlation coefficients to moderate and strong results (Table 2).

**Table 2 -.** Correlation coefficient (R) between the Prevalence of Microcephaly (by Live Birth) and the Density of prevalence of Swine and Bovines for the counties of Rio Grande do Norte, Ceará and Paraíba.

| Variável | R | p value |
| --- | --- | --- |
| *Density of*<br>*Prevalence of*<br>*Swine RN* | 0,5040 | < 0,0001 |
| *Density of*<br>*Prevalence of*<br>*Bovines RN* | 0,5841 | < 0,0001 |
| *Density of*<br>*Prevalence of*<br>*Swine CE* | 0,5281 | < 0,0001 |
| *Density of*<br>*Prevalence of*<br>*Bovines CE* | 0,8063 | < 0,0001 |
| *Density of*<br>*Prevalence of*<br>*Swine PB* | 0,5367 | < 0,0001 |
| *Density of*<br>*Prevalence of*<br>*Bovines PB* | 0,3696 | < 0,0001 |
| *Density of*<br>*Prevalence of*<br>*Swine*<br>*(RN, PB e CE)* | 0,4047 | < 0,0001 |
| *Density of*<br>*Prevalence of*<br>*Bovines*<br>*(RN, PB e CE)* | 0,5235 | < 0,0001 |

In association and concomitantly, the sensitization of the calculation, the homogenization of the results and the biological logic that human exposure to animals is directly related to its density in the territory, could hardly occur by chance (Tables 1 and 2).

## 9. DISCUSSION

This work was motivated by the perception that in Rio Grande do Norte (RN) there was an overlap between the cartographic distribution of microcephaly and the basins of the rivers (Figure 2 and 3) saving the driest regions where, curiously, there were higher rates of infestation by Aedes aegypti [22-24] (Figure 4).

The rivers in the Brazilian Northeast region are surrounded by agriculture and livestock, for this reason the work tab hypothesis was that there could be some association between livestock and the current epidemic of microcephaly.

In Indonesia some studies identified the Zika Virus in farm animals, without clinical disease, suggesting that they could be the reservoir of a silent virus, hard to be detected [26]. ZKV is also able to increase placental permeability [6, 9, 13], which enables fetal aggression, by itself, or by some other associated agent.

The involvement of livestock in the epidemiology of microcephaly may point to BVDV, found in cattle and pigs and endowed with proven potential teratogenic in animals. It is a flavivirus that causes the most prevalent infectious disease of cattle, the bovine diarrheal disease [27]. It can be transmitted vertically and produce malformations and microcephaly in calves [27-34]. Its horizontal transmission is done by body fluids, including milk, saliva and fomites [27-34]. Some studies have confirmed its presence in outbreaks of diarrhea in children [33]. Nevertheless, BVDV has been seen as of little relevance to human pathology [14].

The pigs also become ill by BVDV [35-37]. A recent article identified the prevalence of this virus in 45% of pig herds in the county of Mossoro and in 4% of the animals [37].

The largest prevalence of microcephaly in small towns (Figure 1) discloses a rural environment that is consistent with the involvement of livestock in its epidemilogy and escape from the dynamics of transmission of arbovirus by Aedes aegypti [38, 39]. Studies on the spread of dengue showed that the population density of large cities and the intensity of infestation by the Aedes aegypti peridomiciliary are the two most relevant factors for the amplitude of spread and, combined, produce large epidemics in the larger cities [38,39].

As any complication is always the fraction of a larger series of patients, it would be expected that in the big cities there would be not only the highest number of cases of microcephaly, but also the highest prevalence, which does not occur (Figure 1).

The greatest strength of the correlation coefficient between the Prevalence of microcephaly and the variables that measured the “density of prevalence” of swine and bovines (Tables 1 and 2) does not invalidate, obviously, the hypothesis of participation of Aedes aegypti/ZKV in the epidemiology of microcephaly, but it seems to condition it to the concentration of animals in each territory.

Sensitization of the results of the associations, stronger for the “density” variables, seems to be due to a better sizing of the influence of the concentration of animals as a contagion factor, (Tables 1 and 2) which is consistent with a disease transmitted by a territorial mosquito and / or possibly by fomites, if some involvement of BVDV was proven. The homogenization of the results, especially with the strengthening of the correlation coefficients, as seen in Table 2, means that the measured variables have had, everywhere, a similar behavior, a fact which gives them verisimilitude.

Cases of microcephaly born in counties with more than 100,000 inhabitants and excluded from the analysis (according to the methodology) can be justified by the contagion through milk and unsterilized meat; foods where there is proven survivability of BVDV [26]. This indirect transmission, possibly exacerbated during the period of the epidemic, is linked to consumption and not to physical proximity to livestock, which is the object of this work. These cases always produced prevalences lower than those found in smaller cities.

It is useful to note that the gap data of the agricultural census (ten years) and of the population projection (four years) could mean more robust interaction in the real world than in numbers. The overlap of confirmed cases with the river basins of RN [22, 23] (Figure 2 and 3) and the non overlap of microcephaly with peridomiciliary infestation for that state [24] (figure 4) add biological logic to the livestock involvement hypothesis in the epidemiology of microcephaly, reducing the chances that the associations found are casual. There also did not occur an overlap of the Prevalence of microcephaly with the spatial distribution of the dengue epidemic [25] (Figure 2 and 5), which should have acted as a good marker for the occurrence of other arboviruses.

These results must be comprehended in light of the fact that the epidemiology of this disease is not fully understood. It is important to note that not all the pregnant women affected by Zika in the first trimester of pregnancy had microcephalic or malformed babies [6-8], that the cases of Zika related microcephaly are more frequent in Brazil than in others affected countries [10] and that inside the country the disease is more prevalent in the Northeast region and in low-income families [1, 12]. This set of asymmetries makes ZKV apparently an etiologic agent dependent on another factor to express its most harmful complication, [7, 8, 12].

Linear regressions are not able to define causalities but associations. The present study found correlations ranging from moderate to strong between the prevalence of microcephaly and the variables that measured the concentration of animals in the territory (Table 2). New epidemiological, pathophysiological and etiological studies of the possible connection between livestock and microcephaly are, therefore, mandatory.

